# Visual field advantage: Redefined by training?

**DOI:** 10.1101/476614

**Authors:** Scott A. Stone, Jared Baker, Rob Olsen, Robbin Gibb, Jon Doan, Joshua Hoetmer, Claudia L.R. Gonzalez

## Abstract

In 1990, Fred Previc postulated that most peri-personal space interactions occurred in the lower visual field (LVF), leading to an advantage when compared to the upper visual field (UVF). It is not clear if extensive practice can affect the difference between interactions in the LVF/UVF. We tested male and female basketball varsity athletes and non-athletes on a DynaVision D2 visuomotor reaction task. We recruited basketball players because in their training they spend significant amount of time processing upper visual field information. We found a lower visual field advantage in all participants, but this advantage was significantly reduced in the athletes. The results suggest that training can be a powerful modulator of visuomotor function.

## Introduction

Most of our interactions with the world generally happen in the space just in front of us (peri-personal) or just below us. For example, when eating, writing, reading, cooking, or picking up objects from a surface we are engaging our visuomotor system in the lower visual field (LVF). Importantly, there is evidence that the retina is organized to better support processing of information in the LVF versus the upper visual field (UVF; [1]). Curcio [1] showed that within the peripheral retina, the density of superior hemi-retina ganglion cells (i.e. the part of the retina processing LVF information) is significantly higher than the inferior hemi-retina ganglion cell density processing UVF information. It is possible that this LVF advantage may be the result of evolutionary pressures selecting for foraging and feeding behaviour [2]. Therefore, it is reasonable to expect behavioural differences in visual fields, with LVF being processed more efficiently than UVF.

In fact, studies have demonstrated that humans are more efficient when interacting with objects in the LVF compared to the UVF [3–8]. For example, Danckert and Goodale (2001a) showed that visually guided pointing movements in the LVF are significantly faster and more accurate than equivalent movements in UVF. Similarly, Brown, Halpert [9] showed that grasping behaviours in the LVF performed similarly; they were faster and more accurate than in the UVF. Taken together, these studies are consistent with the theory that the LVF is specialized for processing visual information relevant for action in peri-personal space [10, 11]. Functional magnetic resonance imaging studies have also demonstrated differences in visual field processing [7, 8]. In these studies, participants were presented with objects in either the LVF or UVF and then asked to either perform a reach-to-grasp movement towards the object or simply passively view it. These studies demonstrated greater BOLD activation in the dorsal visual stream, as well as the superior parieto-occipital cortex (SPOC), and the precuneus during LVF reach-to-grasp actions.

In the current study we explore the possibility that visual field differences can be modified with experience (i.e. are they plastic?). It has been suggested that throughout the lifespan, plasticity occurs all over the brain, including visual areas and pathways [12, 13]. We wondered if sports that require a greater amount of attention in the UVF, such as basketball, badminton, or volleyball would reduce the LVF advantage. These sports necessarily require its participants to be trained to attend and respond to UVF. As such it is possible that performance between the LVF and UVF is similar for these athletes. We tested this hypothesis in collegiate-level basketball players, a population trained in UVF performance and compared their behaviour to age and sex matched non-athletes. We used a DynaVision D2 visuomotor training device to assess the movement time of male/female basketball players (athletes) and male/female controls (non-athletes) during a reaction-time task. We predicted a LVF advantage in the control group but no such advantage in the athletes.

## Methods

In this study, 40 right-handed young adults (20 female) participated (mean age: 20 years, sd: 2.24). Both male and female groups consisted of 10 athletes and 10 non-athletes (control). All participants provided written informed consent prior to beginning the experiment. The study was approved by the University of Lethbridge Human Subject Research Committee under research protocol #2015-013.

### Procedure

#### DynaVision D2 Movement Time Task

A DynaVision D2 light board (DynaVision International, USA) (Figure 1) was used to assess movement time. The apparatus consists of a board on which buttons are arranged in concentric rings. Each button contains a light-emitting diode (LED), which can be lit up to elicit a response from the participant (i.e. hit the button). When a button is pressed by the user, the board measures reaction time to the nearest 1/100 of a second. The DynaVision D2 is typically used for athletic training and assessment [14, 15]. Each participant removed their shoes to control for the degree of shoe comfort, and the lights were dimmed to increase the contrast of the LED on the buttons. The board contains a small LCD screen slightly above the center, which was covered so it would not distract the participant (See Figure 1). A white fixation cross, made of tape, was placed in the middle of the board. The board was then adjusted to the participant’s eye-level to evenly split the UVF and LVF. Before starting, we ensured the participant could reach all buttons. The outermost light-ring was deactivated because not all participants could easily reach it. A custom program was created that made a single button light up in a pseudo-random location in either the UVF or LVF, which would change when the participant hit it. Each session lasted for a total of 60 seconds.

**Figure 1.**
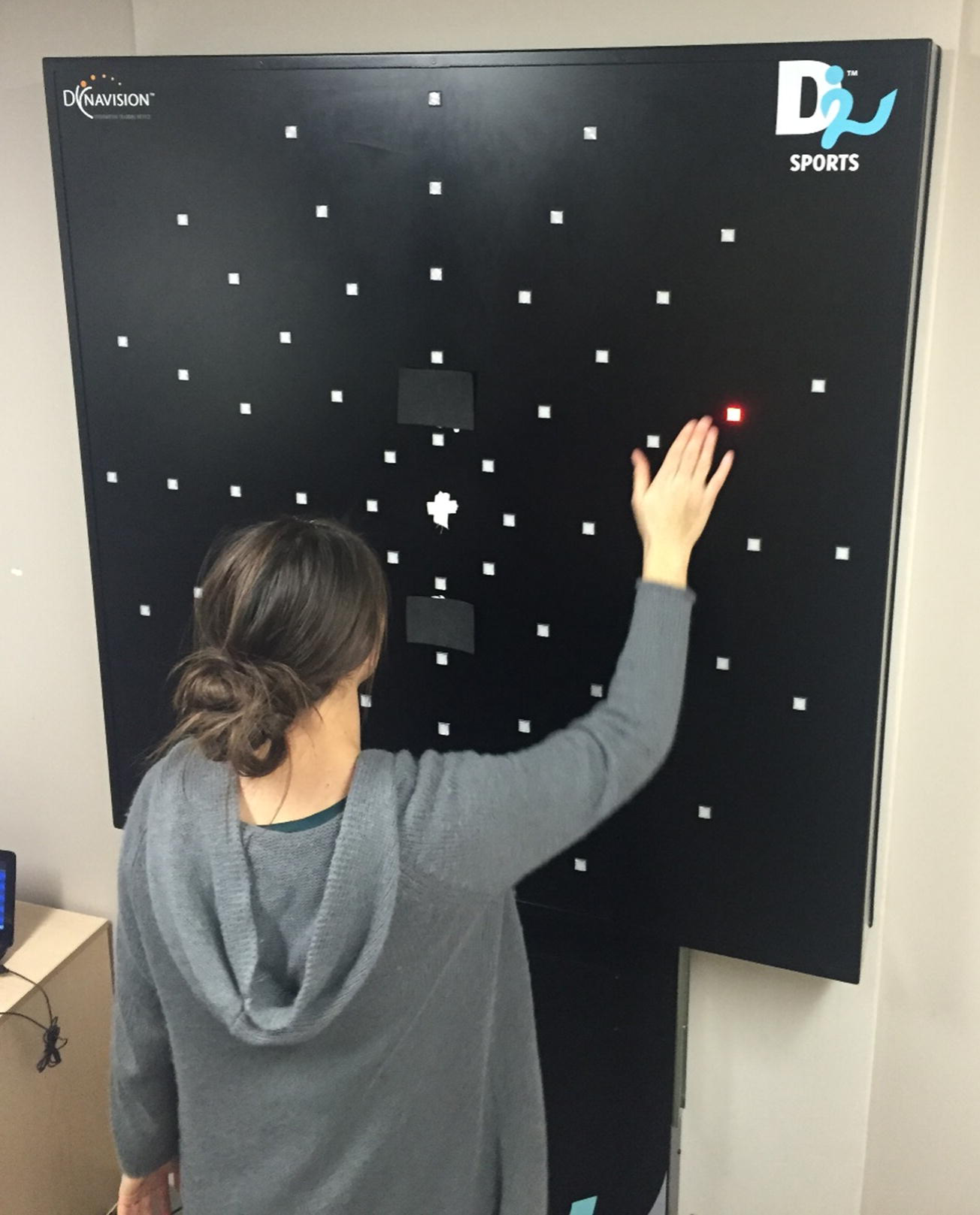
A participant completing the DynaVision reaction time task. The participant fixated on the white cross in the center for the duration of the 60-second session. A single button would light up until the participant hit it. Reaction time for each button press was recorded. Both written and informed consent was obtained from participant for the publication of this image.

#### Movement time & score

The movement time and score were recorded for each session. The average movement time was recorded for each quadrant of buttons. The score was calculated as the total number of buttons pressed during each 60-second session.

#### Practice sessions

To become familiarized with the board, each participant was given two practice sessions lasting 60 seconds each. During these sessions, the lights could appear at any position on the board. Upon confirming the participant understood and was comfortable with the goal of the task, the session would begin. Practice session data was not used in analysis.

#### Sessions

Each participant completed a total of four sessions of 60 seconds each. Two sessions took place in the UVF and the remaining two took place in the LVF. The starting visual field was counterbalanced across participants to eliminate any influence of starting visual field.

#### Trials

The number of trials per session was dependent on the speed of the participant. One trial was equal to one button press. The inter-stimulus-interval was zero as upon pressing the button, a different button would light up. Each button would stay lit until it was pressed.

#### Statistics

The average movement time and score was recorded for each session. All statistical tests were performed on the average of the two sessions in each visual field. Results were considered significant at a p-value below 0.05. All data was analyzed offline using SPSS Statistics 24.0 for Windows (SPSS Inc., Chicago, IL, USA).

## Results

### Handedness questionnaire

All participants self-reported as right-handed. This was confirmed using a Modified Edinburgh Waterloo Handedness Questionnaire [16, 17]. The average score was +32.68(SD: ±2.88) with a possible score in the range of +44 (extremely right-handed) / −44 (extremely left-handed).

### DynaVision

#### Movement time – UVF versus LVF

The movement time for each button press was calculated as the time between the button first lighting up and being pressed. A repeated-measures ANOVA with visual field (upper/lower) as within factors and athletic status (athlete, non-athlete) and sex (female, male) as between factors was conducted. The results showed a main effect of UVF/LVF (F(1,36) = 68.15; p < 0.0001, η2 = 0.654), a main effect of athletic status (F(1,36) = 22.16; p < 0.0001, η2 = 0.381), but no main effect of sex (F(1,36) = 2.48; p = 0.12, η2 = 0.064). Participants responded faster in the LVF (mean = 619ms; sd: 131ms, se: 20ms) when compared to the UVF visual field (692ms; sd: 91ms, se: 14ms). Athletes (mean = 592ms; sd: 49ms, se: 11ms) were faster in their responses than non-athletes (mean = 720ms; sd: 130ms, se: 30ms). Importantly, there was a significant interaction (Figure 2a) between UVF/LVF and athletic status (F(1,36) = 16.46; p < 0.0001, η2 = 0.314). Although participants in both groups reacted faster to stimuli in the LVF, the difference was greater in the non-athletes group (Athletes: (t(19) = 4.25; Non-Athletes: (t(19) = 7.02). No other interactions were significant (p > 0.05).

**Figure 2.**
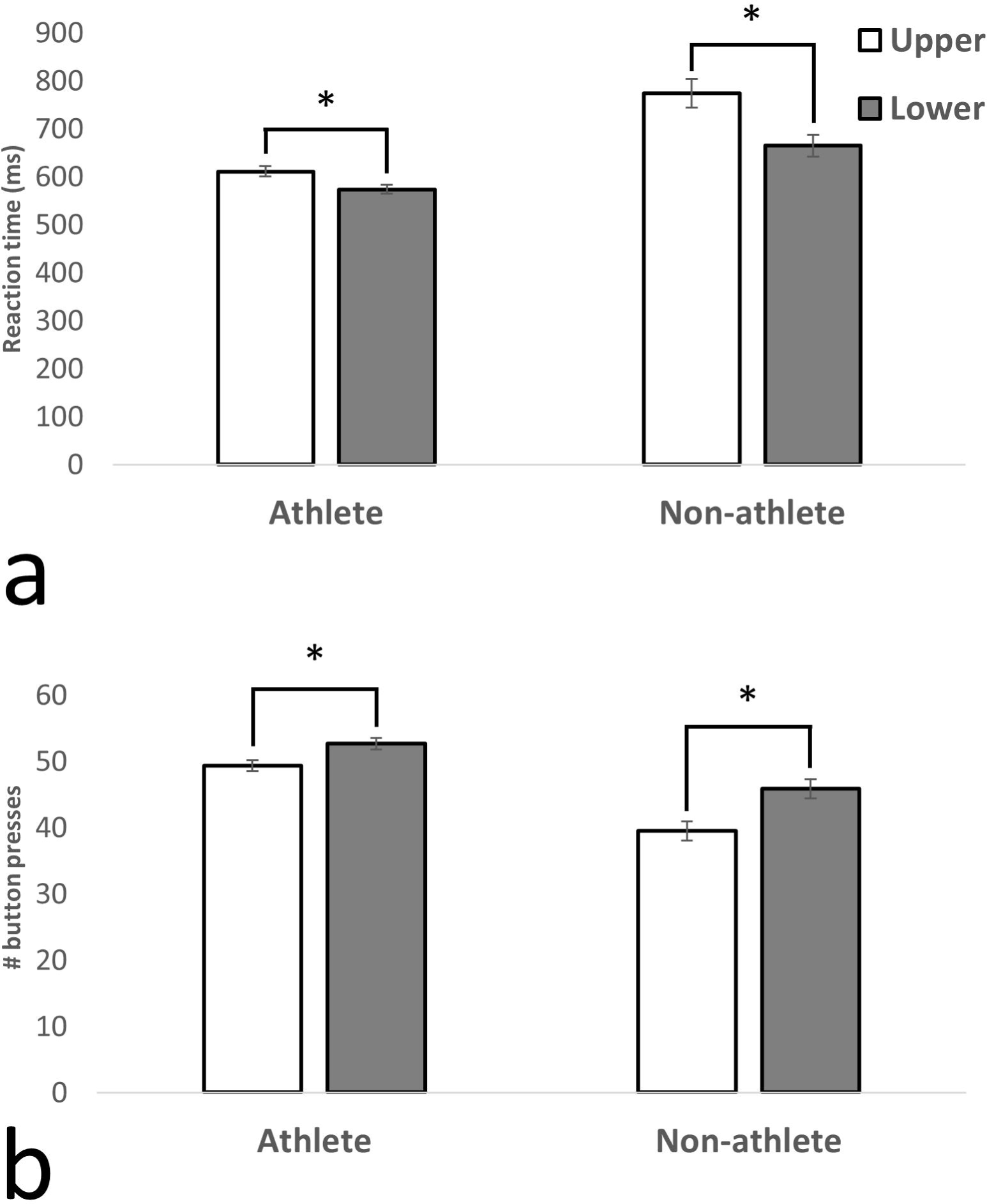
A bar graph illustrating the average MT (A) and number of buttons pressed (B) in the LVF and UVF within the athlete or non-athlete groups. A significant main effect of visual field was revealed, with participants’ LVF responses being faster than UVF, regardless of athletic status. Standard error of each measure is shown. A significant interaction between athletic status and visual field was revealed. The differences in MT and number buttons pressed between the visual fields were smaller in athletes than in non-athletes.

#### Scores – UVF versus LVF

The score was calculated as the number of buttons correctly hit by the participant during the 60 second session. A repeated-measures ANOVA with visual field (upper/lower) as within factors and athletic status (athlete, non-athlete) and sex (female, male) as between factors was conducted. The results showed a main effect of UVF/LVF (F(1,36) = 73.68, p < 0.0001, η2 = 0.672), a main effect of athletic status (F(1,36) = 28.14, p < 0.0001, η2 = 0.439) (Figure 2b.), but no main effect of sex (F(1,36) = 2.98, p = 0.093, η2 = 0.076). Participants hit more buttons in the LVF (mean = 49.2; sd: 6.37, se: 1.42) when compared to UVF (mean = 44.4; sd: 7.19, se: 1.60). Athletes (mean = 50.97; sd: 3.50, se: 0.78) hit more buttons than non-athletes (mean = 42.6; sd: 6.21, se: 1.38). Similar to the results of movement time, a significant interaction (Figure 2b) was detected between UVF/LVF and athletic status (F(1,36) = 7.12, p < 0.05). There was a significant difference between the number of buttons successfully hit in the UVF and LVF in both groups, but the difference was greater in the non-athletes group (Athletes: (t(19) = −4.66); Non-Athletes: (t(19) = −7.33). No other interactions were significant (p > 0.05).

#### Movement time – left versus right VF

Although our main question focused on differences between the LVF and UVF we conducted similar analyses on the left/right visual fields for movement time and score. None of the analyses (main effects or interactions) were significant (all *Ps* > 0.3).

## Discussion

The present study had two investigative goals: 1) To quantify the MT difference within the UVF versus LVF using the DynaVision D2 basic visuomotor movement time task. 2) To determine if athletes, specifically basketball players, display experience-dependent plasticity in the UVF. Results showed that basketball players were faster than controls. In addition, all participants had consistently lower RTs in the LVF as compared to the UVF. Further, a significant interaction between the visual field (UVF/LVF) and athletic status (i.e. varsity basketball player or control) was revealed (Figure 2). The difference in MT between the UVF and LVF was reduced in the basketball players. This suggests that the experience the athletes had in their basketball training quickened their MT in the UVF. While it is possible the differences observed are due to the athletes’ biomechanical advantages, this is unlikely because no differences were discovered when comparing the left and right visual fields. This suggests that the differences were 1) specific to the upper and lower visual fields and 2) due to a visuomotor coupling advantage only present in the athletic group.

Overall, we found that the athletes were faster than the non-athletes, which might be expected due to the structured training regimens adhered to by the athletes [18]. Allard and Starkes [18] recruited volleyball players and non-athletes to complete a task where the goal was to detect a volleyball in a rapidly presented slide. They found that while accuracy was similar between the groups, the volleyball players were significantly faster than their non-athletic counterparts. Furthermore, greater breadth of attention was reported in elite athletes when compared to novices, and that such differences varied as a function of athletic expertise [19]. In this study, the ability to devote attention to different objects was quantified as a function of athletic expertise. For example, soccer players were found to perform better at tasks that require greater horizontal breadth of attention whereas volleyball players show a similar effect in vertical space. These results align with the findings of the current study.

Studies investigating visual fields for differences have demonstrated increased efficiency in the LVF for visuomotor processing [3, 4, 6–8]. In the Danckert and Goodale (2001) experiment, a pointing task was used to demonstrate that responses to targets in the LVF were always faster than in the UVF. Furthermore, as target size decreased, movement time and accuracy increased but only in the LVF. In other words, target size processing in LVF appears to be more sensitive. In contrast, movement time in the UVF does not seem to correspond to target size, suggesting less attention is given. The authors suggest that this is due to LVF’s natural superiority in processing visual feedback, where the LVF has a functional bias for these types of movements. This result makes sense in the context of Curcio’s (1990) finding of higher ganglion cell density in the peripheral retina that processes LVF information as compared to the UVF. This implies the LVF information is processed more efficiently, even pre-cortically. The results of the present study agree with Danckert and Goodale, as the LVF movement times were consistently lower than UVF. We suggest the lower movement times observed in LVF are driven by the functional bias of LVF for this type of stimulus.

It is possible that UVF indeed requires more effort to interact with, on both a muscular and visual processing level. Given that males in general tend to have significantly more muscle mass in the upper body than females (e.g. Janssen et al, 2000), we would expect to find significant differences between males and females for this task. As we do not find any difference in any of the measures, this effect is not likely simply driven by muscle mass differences in the groups. While the basketball players likely do have increased muscle mass in the upper body, it is unlikely it is significantly changing their performance in the UVF versus LVF portions of the board. While it is difficult to directly measure the influence of neuroplasticity as a result of visual system training, we feel that this is an appropriate ecologically valid task to assess this measure. Given that it is indeed harder to interact in the UVF, it makes ethological sense that the upper visual field would be under-represented in attention. Extensive training would enhance function in this area and result in better performance in those who trained more (i.e. basketball players).

The plastic nature of the brain allows for dynamic reorganization [13], especially when paired with endurance training regimens such as those used by varsity sports teams [20–22]. We specifically recruited varsity basketball players as our athletic group because of the increased demand and exposure to UVF processing. Zwierko, Lubiński [23] measured visual evoked potentials (VEPs) in female volleyball players just prior to and following two years of intensive training. They found the latency of key visual conductivity signals in the VEP waveform was reduced after the training. Interestingly, they reported that the latency of the N75 (which is thought to originate in the primary visual cortex) was significantly reduced after training for stimuli occurring on the peripheral retina. In essence, training modified visual cortex activity through experience-dependent plasticity initiated at the peripheral retina. This is in line with the results of the current experiment because we propose that the lower movement times observed in the athletes are directly caused by plastic changes initiated at the level of the peripheral retina. We speculated that the increased amount of time basketball players spend processing stimuli in UVF would lead to an increase in performance in that field, ultimately reducing the advantage over LVF. This is precisely what we found; UVF processing was enhanced in the athletic group (relative to non-athletes), resulting in a decreased (yet significant) difference between visual field RTs. It is possible that this enhancement is driven by cortical plasticity in the visual and visuomotor pathways, which continue to change as a result of experience throughout the lifespan [12]. Jensen, Marstrand [24] measured motor evoked potentials (MEPs) during a simple visuomotor task that involved moving the elbow to match patterns shown on a computer screen. The MEPs (measured via transcranial magnetic stimulation to motor cortex) were significantly increased after training, suggesting visuomotor training had affected visual and motor cortex connectivity. It is also worth noting that control of the elbow is performed by proximal muscle groups, which receive less corticospinal control [25] and are thought to be more important when playing most sports. Because basketball players spend a large amount of time processing UVF stimuli (e.g. looking for passes, watching the basketball hoop) and acting on those stimuli through motor coordination, it is reasonable to suppose that better performance in this field results from practice. Neuroimaging studies are needed to evaluate this speculation.

One final consideration is the lack of sex differences; we did not find a significant main effect of sex on MT in either visual field nor a significant interaction. Although some studies have found differences between the sexes in visuospatial tasks, it is possible that the difference in processing abilities between the UVF and LVF are so robustly conserved that sex has no effect on performance for this task.

In conclusion, we created a task and methodology to measure whether or not training and experience could change the typical performance difference between LVF- and UVF-processing. The results demonstrated this to be the case, suggesting that even the highly conserved differences in information processing in LVF and UVF can be modified through experience. The current finding has implications for both training and rehabilitation after nervous system damage.

